# Detection of Chemotherapy-Resistant Pancreatic Cancer Using a Glycan Biomarker

**DOI:** 10.1101/2020.06.01.128082

**Authors:** ChongFeng Gao, Luke Wisniewski, Ying Liu, Ben Staal, Ian Beddows, Dennis Plenker, Mohammed Aldakkak, Johnathan Hall, Daniel Barnett, Mirna Kheir Gouda, Peter Allen, Richard Drake, Amer Zureikat, Ying Huang, Douglas Evans, Aatur Singhi, Randall E. Brand, David A. Tuveson, Susan Tsai, Brian B. Haab

**Author notes:** Corresponding Author Brian B. Haab, PhD, Van Andel Institute, 333 Bostwick NE, Grand Rapids, MI 49503. Author Contributions B.H. and C.G. conceived and designed the experiments. C.G., L.W., Y.L., and B.S. conducted most of the laboratory experiments. J.H. and D.B. assisted in data analysis. I.B. analyzed the RNA-seq data and performed bioinformatic analysis. D.P and D.A.T generated organoids slides and conditional media. H.Y. performed statistical analyses and assisted in data interpretation. M.A. and S.C provided patient serum samples and contribute to data analysis. M.K.G., P.A., R.D., A.Z., D.E., A.S., and R.E.B provided specimen resources and assisted in data interpretation. B.H. wrote the manuscript, and all authors reviewed and commented on the manuscript.

## Abstract

**Background and Aims:** A subset of pancreatic ductal adenocarcinomas (PDACs) is highly resistant to systemic chemotherapy, but no markers are available in clinical settings to identify this subset. We hypothesized that chemotherapy-resistant PDACs express a glycan biomarker called sTRA. *Methods*. We tested this marker to identify treatment-resistant PDAC in multiple systems: sets of cell lines, organoids, and isogenic cell lines; primary tumors; and blood plasma from cohorts of human subjects. *Results*. Among a panel of 27 cell lines, high levels of cell-surface sTRA identified higher resistance to seven chemotherapeutics used against PDAC. Using primary tumors from two different cohorts, patients who were positive for a gene-expression classifier for sTRA received no statistically significant benefit from adjuvant chemotherapy, in contrast to those negative for the signature. In another cohort, using direct measurements of sTRA in tissue microarrays by quantitative immunofluorescence, patients who were high in sTRA again had no statistically significant benefit from adjuvant chemotherapy. Further, a blood-plasma test for the sTRA glycan identified the PDACs that showed rapid relapse following neoadjuvant chemotherapy. This blood test performed with 96% specificity and 56% sensitivity in a blinded cohort using samples collected before the start of treatment. *Conclusion*. These findings establish that tissue or plasma sTRA can identify PDACs that are resistant to neoadjuvant or adjuvant chemotherapy. This capability could help apply systemic treatments more precisely and facilitate biomarker-guided trials targeting resistant PDAC.

## Introduction

Systemic therapy is considered necessary for all patients with pancreatic ductal adenocarcinoma (PDAC), even those with localized disease, because most patients already have occult metastases at the time of diagnosis^1^. Chemotherapy particularly benefits patients who have surgical resection of the tumor. In a seminal study that established adjuvant chemotherapy (chemotherapy applied after surgery) as standard of care for PDAC, the median disease-free survival after surgery improved to 13.4 months with gemcitabine from 6.9 months with observation^2^. Further improvements in chemotherapy were demonstrated using the stronger FOLFIRINOX regimen^3^, or gemcitabine in combination with capecitabine^4^ or nab-paclitaxel^5^. Systemic therapy is increasingly applied prior to surgery—called neoadjuvant therapy—in order to increase the percentage of patients who receive chemotherapy^6^, since some patients have a delay or reduction in their adjuvant chemotherapy as a consequence of surgery. Neoadjuvant therapy could have the additional advantages of identifying patients with rapid progression who would not benefit from surgical intervention; treating occult metastases earlier; and downsizing the tumors to increase the chance for a margin-free (R0) resection^6^.

While the combination of surgery plus systemic therapy results in significant benefit relative to surgery alone, a subset of PDACs is highly resistant to systemic therapy. Nearly 40% of patients receiving surgery plus gemcitabine monotherapy experience relapse within one year of surgery^2^. Even in the subset of fit patients who are candidates for more aggressive chemotherapy regimens, over 25% relapse within one year^3^. Currently, identifying this chemotherapy-resistant cohort prior to treatment remains a challenge, since conventional imaging, liquid biopsy, and molecular biomarkers are lacking.

The gene-expression subtypes defined in prior research^7-9^ potentially provide some guidance to this problem. The consensus subtypes have been termed classical, basal (also referred to as quasi-mesenchymal), and exocrine, terms chosen to reflect the normal cell types that most closely correspond to the cancer cells. In retrospective evaluations of outcomes following curative resection, tumors with transcriptome profiles matching with the classical subtype had longer survival than the others^7-9^. Likewise, among patients with metastatic PDAC, the classical subtype was associated with longer survival in retrospective analyses^10, 11^. On the other hand, patients with the classical subtype demonstrated no benefit from adjuvant chemotherapy^9, 12^, in contrast to patients with the basal type, and cell lines of the classical subtype are more resistant to chemotherapy than those of the basal type^8^. Therefore, the predictive role of molecular subtyping in PDAC treatment remains to be established.

Further exploration of the role of molecular subtyping in PDAC requires a practical biomarker. Glycans (which are oligosaccharides) are an intriguing option, given their abundance on cell surfaces and secreted proteins, and because specific structures are indicators of cell differentiation. Glycans altered in pancreatic cancer include CA19-9^13^, members of the Lewis blood group family^14^, and ABO blood group antigens^14^.

We recently identified a new biomarker of PDAC that is a cell-surface and secreted glycan called sTRA (sialylated tumor-related antigen)^15, 16^. The glycan is elevated in about half of the cases that do not show elevated CA19-9. A combination blood test using CA19-9 and the new marker performed better than CA19-9 alone in blinded tests for distinguishing PDAC from benign diseases^17^. The tumors expressing primarily sTRA tended to be sparse, poorly differentiated, or highly vacuolated, while those expressing mainly CA19-9 were part of well differentiated or moderately differentiated secretory glands^16^. These facts suggested that the two groups represent distinct subtypes of tumors having differing biology and clinical behavior. Of particular interest was the possibility that the drug-resistant group observed in clinical care corresponds to a glycan-defined subtype. We explored that possibility using multiple model systems and cohorts of clinical specimens.

## Materials and Methods

### Cell Culture and Reagents

The PaTu-8988S and PaTu8988T cell lines were obtained from Creative Bioarray (Shirley, NY), and Colo357, L3.3, and L3.6PL lines were kindly provided by Dr. Isaiah J. Fidler (University of Texas, MD Anderson Cancer Center). The remaining cell lines were obtained from ATCC (Manassas, VA). All cell lines were cultured in RPMI-1640 supplemented with 5% fetal bovine serum, 2 mM L-Glutamine, and 100 IU/mL penicillin/streptomycin. The cells were grown at 37 °C in a humidified atmosphere supplemented with 5% (v/v) CO_2_. The chemotherapeutic reagents cisplatin, etoposide, gemcitabine, and 5-FU were obtained from Sigma (St. Louis, MO). Irinotecan, oxaliplatin, and paclitaxel were obtained from Cayman Chemical (Ann Arbor, MI). All drugs were dissolved in dimethyl sulfoxide or dimethylformamide. For the preparation of FOLFIRINOX, 5□-FU was first prepared in dimethylformamide, and the leucovorin, irinotecan, and oxaliplatin were each added in a 1:5 ratio by weight to 5□-FU. The concentration of FOLFIRINOX was calculated based on the 5□-FU concentration.

### Drug Treatment Studies

Cells were seeded into 96 well plates at 2 × 10^3^ cells per well and cultured for 3 d before treatment with a drug or drug mixture at six different concentrations each. After 3 d, cell viability was estimated using CellTiter-Glo (Promega, Madison, WI). The IC_50_ values were calculated using GraphPad Prism 6 (GraphPad Software, San Diego, CA) with 5-parameter, variable-slope fits.

For the outgrowth of drug-resistant L3.3 sublines, the cells were cultured in medium containing cisplatin (2 *µ*M), etoposide (2 *µ*M), gemcitabine (0.02 *µ*M), or 5-FU (5 *µ*M) for 3 d and recovered for 4 d in medium without drug. This process was repeated for three rounds, after which the cells were maintained in a 1/10 of concentration of the drug used for selection.

### Statistics

Differences between marker-positive and marker-negative cell lines in IC_50_ and percent viability were tested using the Mann-Whitney U test. Differential expression in the RNAseq data was tested using empirical Bayes quasi-likelihood F-tests, and the p values were adjusted using the Benjamini-Hochberg method. Differences in OS between patient groups in the survival analyses were evaluated with the log-rank test. Differences between patient groups in proportions of patients with long or short OS were analyzed with the Fisher’s Exact test and the Breslow-Day test for homogeneity of odds ratios. Differences in sensitivity and the average of sensitivity and specificity were analyzed with the Wald test based on bootstrap SE estimate. P values of less than 0.05 were considered significant.

To perform cross validation with bootstrapping, the patients were randomly divided into five groups stratified on case-control status. Four groups served as the training subset to derive the panel and one group served as the testing subset to estimate the panel performance. The five estimates of performance were then averaged to get the final performance estimate. This cross-validation was repeated on 200 samplings of the cohort to compute the standard error for the cross-validated performance estimate.

### Human Specimens

The plasma samples were collected under protocols approved by the Institutional Review Boards at the University of Pittsburgh Medical Center and the Medical College of Wisconsin. The tissue samples for tissue microarrays were collected under approved protocols at the Medical University of South Carolina, University of Pittsburgh Medical Center, and Memorial Sloan Kettering Cancer Center. All subjects provided written, informed consent, and all methods were performed in accordance with an assurance filed with and approved by the U.S. Department of Health and Human Services.

The plasma collections took place prior to any surgical, diagnostic, or medical procedures. The donors consisted of patients with pancreatic cancer or a benign condition involving the pancreas, and from healthy subjects. All blood samples (EDTA plasma) were collected according to the standard operating procedure from the Early Detection Research Network and were frozen at −70 °C or colder within 4 h of time of collection. Aliquots were shipped on dry ice and thawed no more than three times prior to analysis. Disease progression was diagnosed radiographically based on CT scans performed at 3-4 month intervals for the first two years and at six month intervals thereafter. Occasionally, a tissue biopsy was performed.

## Results

We initially tested for differences between PDACs classified by glycan expression using a panel of 27 cell lines. We classified each cell line based on the sTRA glycan or the CA19-9 glycan (Figure 1A). Both glycans are capped with sialic acid on type-1 *N*-acetyl-lactosamine (LacNAc), the disaccharide of galactose linked β1,3 to *N*-acetyl-glucosamine(GlcNAc), and the CA19-9 epitope has a fucose attached to the N-acetyl-glucosamine, which is necessary for its recognition by selectin receptors. Type-1 LacNAc, as recognized by the TRA-1-60 antibody^18^, is a marker for induced pluripotent stem cells, but the sialylated version has not been well studied due to lack of an effective antibody. We indirectly detected the sialylated structure using sialidase to uncover the TRA-1-60 epitope (Figure 1A).

**Figure 1.**
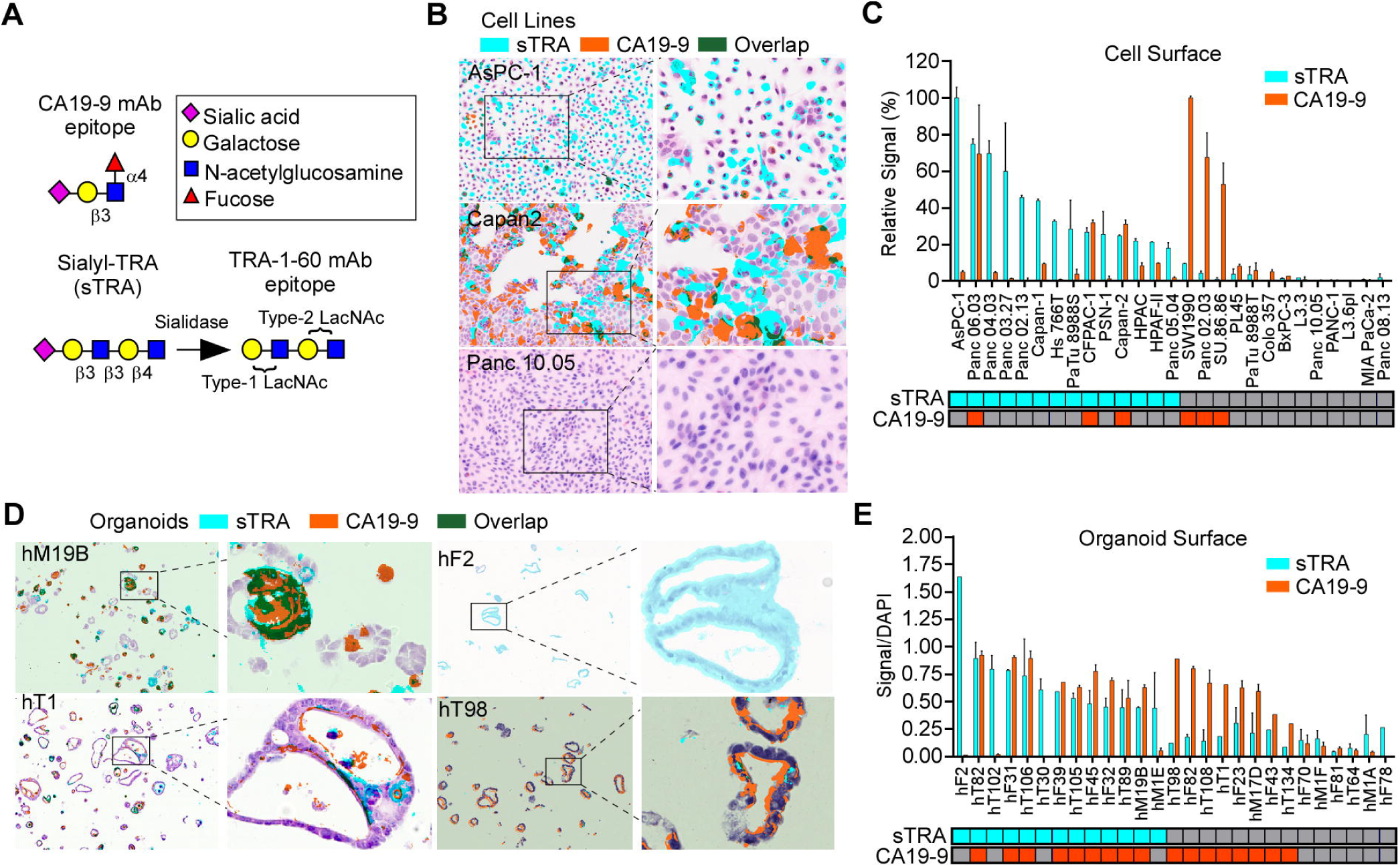
Complementary glycan expression in model systems. A) Glycan structures. B) Diverse sTRA and CA19-9 expression in cell culture. C) Cell-surface expression of 27 cell lines. The classification of positivity at the bottom was based on a cutoff of signal-to-noise ratio > 3. D) Diverse glycan expression in organoid models. E) Cell-surface glycan expression of 27 organoid models. The classification of positivity at the bottom used cutoffs that optimized the discrimination between background signal and true marker expression. The magnification was 20X for the non-zoomed images.

The cell-surface expression of sTRA and CA19-9 was variable among the cell lines, with some primarily expressing only one glycan and others expressing both or neither (Figs. 1B and 1C). Organoid models of pancreatic cancer^19, 20^ likewise showed variable expression of one, both, or neither of the glycans (Figure 1D), with slightly different proportions among them (Figure 1E). The difference in proportions could be due to cellular selection, effects of culture conditions, or merely random variation within the small sample size. Nevertheless, these two model systems agree in showing distinct expression of the glycans.

### Gene-expression programs distinguishing the glycan-defined subtypes

Using the cell lines and organoids, we then asked whether similar differences exist in gene transcription programs. A total of 267 genes were differentially expressed between the sTRA-expressing cell lines (not including the three lines also expressing CA19-9) and all others (Figure 2A and Supplemental Table 1). No individual genes were differentially expressed between the CA19-9 and sTRA groups at p < 0.05 after multiple-hypothesis correction, possibly due to the lower number of CA19-9-positive lines. The sTRA-associated genes had ontologies that were enriched in developmental, drug metabolism, and glycan-biosynthesis pathways (Figure 2A). The developmental gene *BMP4* was a strong individual marker of sTRA cells, as was *CYP3A5* (Figure 2A), a gene previously identified as a marker of PDACs identified as classical and exocrine^21^. In addition, 9 of 14 sTRA-expressing lines were identified as classical, based on the gene classifier called PDAssigner ^8^, compared with 2 of 6 CA19-9-expressing and 0 of 10 glycan-negative lines (Figure 2A), suggesting that sTRA is more likely to recognize the classical subtype. Gene-set enrichment analysis showed that sets defining stem-like differentiation, stem-like metabolism, and the classical subtype were enriched in the sTRA-expressing cells (Figure 2B and Supplemental Table 1). Among individual genes that have been proposed as markers of class, the expressions of *GATA6* and *CYP3A5* were higher in the sTRA cells (Figure 2C). *KRT81* and *HNF1A* showed weak associations with sTRA.

**Figure 2.**
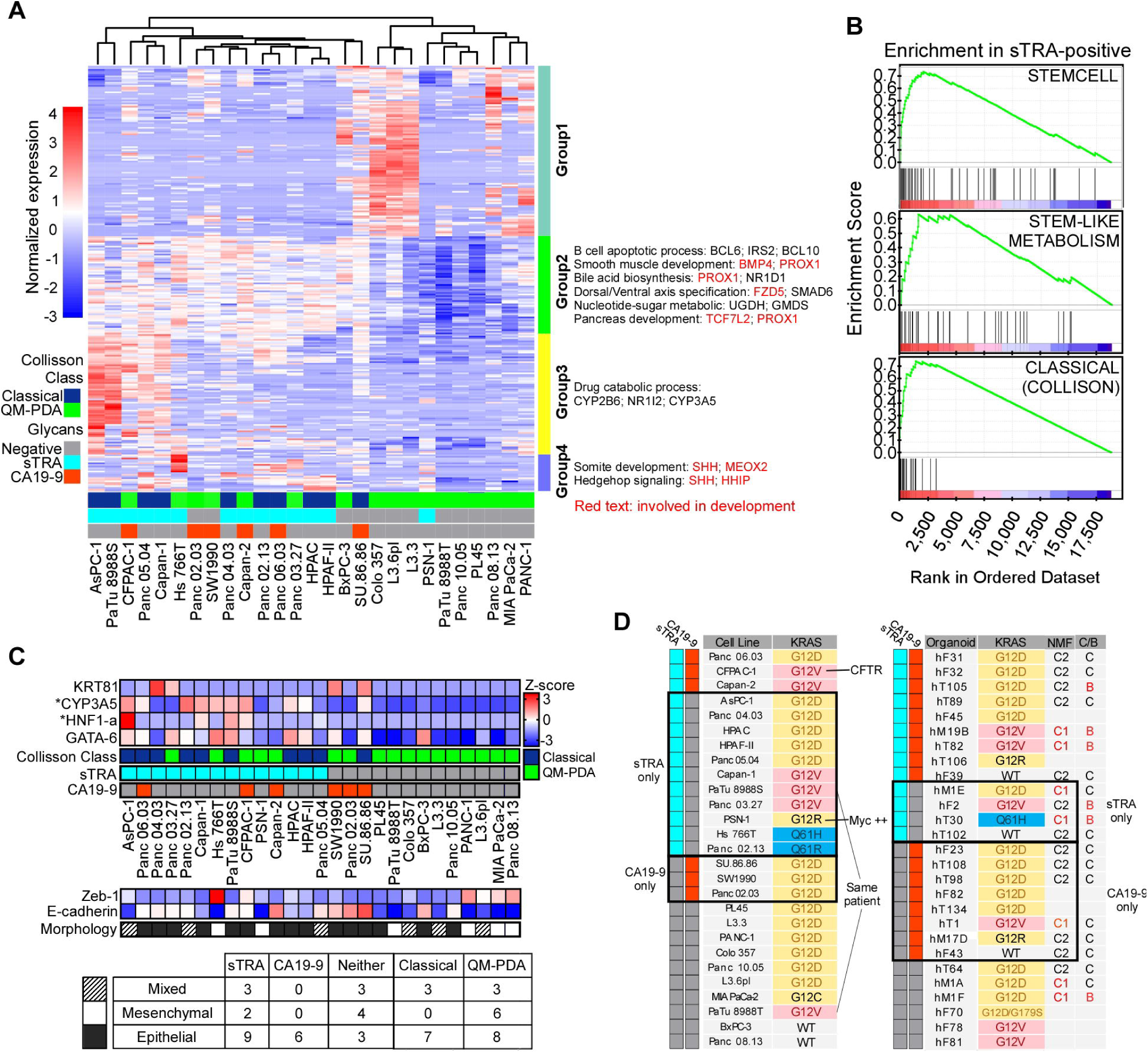
Differential gene expression. A) Differentially expressed genes and pathways. B) Enrichment of previously identified sets in the sTRA-expressing cells. C) Relationships between class markers, morphology, and glycan expression across the cell lines. CYP3A5 and HNF1-a were significantly elevated in the sTRA group compared with the non-sTRA group after multiple-testing correction D) KRAS mutation status associated with the glycan-defined groups. NMF, nonnegative matrix factorization to identify C1 and C2 clusters in the organoids. The C/B column indicates the classical or basal status based on a gene set developed for the organoids.

The epithelial/mesenchymal state of cancer cells has been widely explored as an indicator of their origin, invasiveness, or overall tumor-forming aggressiveness. All six of the CA19-9-expressing cell lines were epithelial, as determined by the gene expression of Zeb-1 and E-cadherin (Figure 2C) and by morphology (Supplemental Figure 1), but the sTRA-expressing cells and those expressing neither glycan were of various types (Figure 2C). The organoid models all had epithelial morphologies, but three of the models had mesenchymal characteristics (high Zeb-1 and low E-cadherin) by gene expression (not shown). Two of these produced sTRA exclusively and the third produced neither glycan. The data from both model systems suggest that some sTRA-expressing cancer cells have the potential for mesenchymal-like differentiation, in contrast to CA19-9-expressing cells.

The organoid models were dissimilar to the cell lines in their overall gene expression profiles. All but one of the organoids had classical subtype expression by PDAssigner. However, the gene sets used to divide the organoids^20^ into C1/C2 and classical/basal groups did not separate the cell lines into similar groups (not shown). This finding could be because all of the organoids were epithelial, or it could derive from differences in culture conditions.

The type of KRAS mutation in cancer potentially can drive differences in phenotype^22^. The less-common Q61 alteration appeared exclusively in the cell lines and organoids that expressed only sTRA (Figure 2D). The G12V mutation was in 3 of 11 cell lines and 1 of 3 organoids that expressed only sTRA, in comparison to 0 of 3 cell lines and 1 of 8 organoids that expressed only CA19-9. The common G12D mutation and the rare wild-type cases were found throughout the subtypes. While these observations are based on relatively small sample sizes, they suggest that the Q61H/R mutation fosters cancers that express sTRA in the absence of CA19-9. Mutations in other cancer-associated genes were not more frequent in either glycan-defined group (not shown).

### Resistance to chemotherapy in sTRA-high cultures

We determined the resistance of the 27 cell lines to eight chemotherapeutics that are either front-line or alternative treatments against pancreatic cancer. In a single-dose study, the sTRA-expressing cell lines were more resistant than the sTRA-negative cell lines in each case (Figure 3A). In dose-response analyses to obtain the IC_50_ concentrations (Figure 3B, 3C, and Supplemental Figure 2), the sTRA cells were significantly more resistant than the non-sTRA cells (p < 0.05, Mann-Whitney U Test) to 6 of the 8 drugs. For gemcitabine, the resistance was higher in the sTRA group but with less statistical significance (p = 0.07, Mann-Whitney U Test); for oxaliplatin, resistance was similar between the groups. In contrast, CA19-9 did not define a resistant group (Supplemental Figure 2). Two outlier lines that expressed sTRA but were sensitive—PSN-1 and CFPAC-1—had mutations that were not consistent with the other sTRA-positive cells lines (Figure 2D).

**Figure 3.**
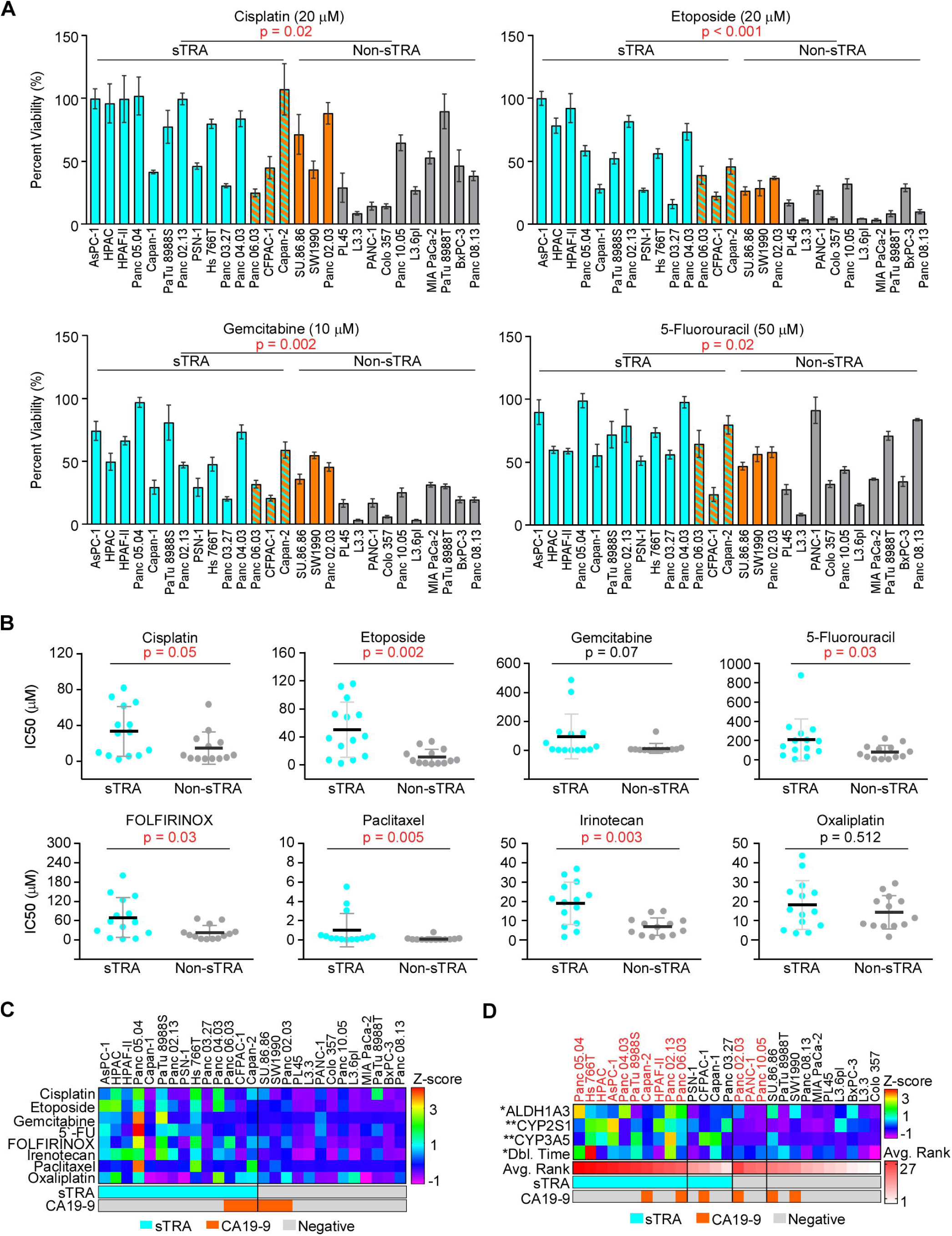
Drug resistance differences between glycan-defined types. A) Single-dose viability in the panel of 27 cell lines. B) IC_50_ values calculated from dose-response curves, grouped by marker group. P Values were calculated by Mann–Whitney test. C) Summarized IC_50_ values for all drugs and cell lines. Z-scores were used to normalize the scales of the IC_50_ values for comparisons. D) Factors and markers associated with resistance. Z-scores were used to normalize the gene-expression data and the doubling times. Avg. Rank is the average across drugs of the ranks in IC_50_ of each cell line among the panel of 27.

We asked whether the high resistance corresponded to traits that have been associated with resistance. The sTRA-positive lines had higher levels of drug-metabolizing enzymes from the cytochrome P450 family (p < 0.05, Mann-Whitney U test), and they had trends toward higher levels of the stem marker ALDH1A3 and longer doubling times (Figure 3D). Thus, some sTRA lines have resistance traits, but the mechanism of resistance may differ between cell lines.

We further tested the above relationships using sets of isogenic cell lines, where all cell lines in a set are from the same individual (Supplement Figure 3). To generate sublines with increased drug resistance, we repeatedly cultured the L3.3 cell line in sublethal concentrations of each of four drugs, followed by recovery and outgrowth of the surviving cells. We found that sTRA expression was increased in several of the sublines, both in the percentage of stained cells and in total staining intensity. Genotyping across over 15,000 SNPs confirmed agreement in genotype between the sublines (not shown). Consistent with the results across diverse cell lines, the sublines with higher sTRA coincided with significantly increased resistance to the drugs.

The cell lines PaTu8988S and PaTu8988T were derived from a single patient and also showed differences in sTRA. The PaTu8988S cell line had much higher sTRA (Supplemental Figure 3) (neither expressed CA19-9), and in accord with the results above, it showed higher resistance to 3 of 4 chemotherapies. These findings from isogenic cell lines parallel the findings from the panel of cell lines and support the idea that sTRA is a biomarker for a subtype of PDAC having high drug resistance.

### Predictive value of the sTRA levels in primary tumors

Next, we tested whether the sTRA levels in primary tumors are associated with resistance to systemic chemotherapy. We determined sTRA levels in two ways, by a gene-expression classifier and by immunofluorescence. To develop a gene-expression classifier, we identified the significantly up-regulated or down-regulated genes (p < 0.02, Bayes quasi-likelihood F-test after multiple-testing correction) associated with sTRA expression in the panel of 27 cell lines (Supplemental Table 1) and used the algorithm from PDAssigner^8^ to assign classes. We used this algorithm because of its previous robust performance and its simplicity for adoption with new gene sets. We applied the classifier to 150 cases of PDAC from The Cancer Genome Atlas^23^ and to 180 cases from the International Cancer Genome Consortium^24^ that had survival information. In both cohorts, distinct groups of patients showed overall differences in expression between genes associated with high sTRA and those associated with low sTRA (Figure 4A). Tests of group differences in central tendency using the classifier genes showed significance (p = 0.001, Adonis test, Vegan R package). This finding confirmed the consistency between the cell lines and both cohorts in the differential expression of the gene groups, and it supports the idea that the classifier identifies true subtypes rather than random variation in expression patterns.

**Figure 4.**
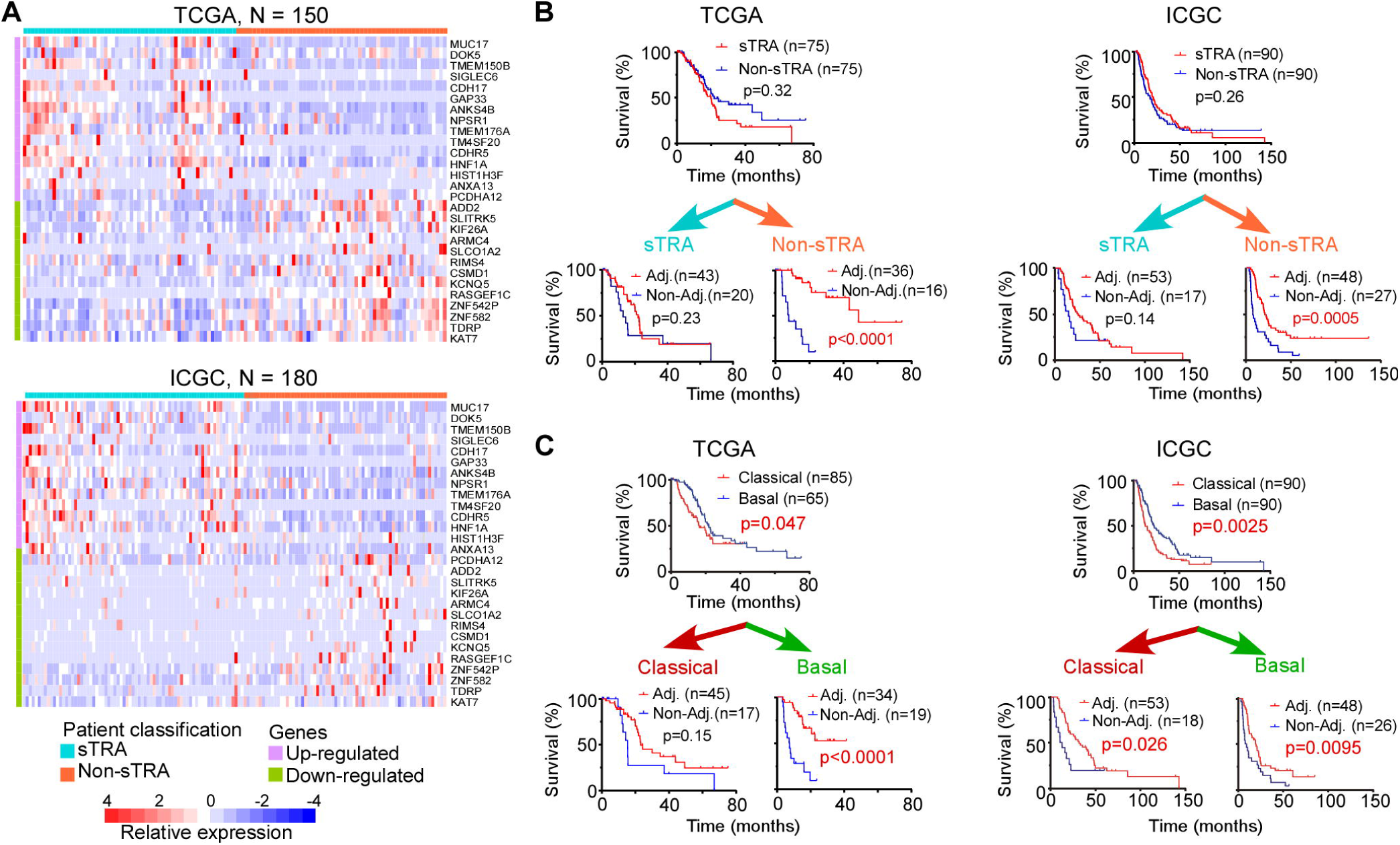
A gene-expression classifier associated with drug resistance. A) Classification of patients using a signature for sTRA. The 28-gene classifier applied to the TCGA (top) and ICGC (bottom) datasets produced two distinct groups of patients. The color bar at top shows the classification used in the subsequent analyses. B) Survival curves grouped by the gene-expression classifier for sTRA positive or negative. C) Survival curves grouped by classical or basal status. The classical and basal classes were from a previous publication for the TCGA dataset and were calculated from the data for the ICGC dataset. The p values are based on the log-rank test.

We assigned the patients to an sTRA or non-sTRA group based on the median of the calculated score of the classifier. No difference in overall survival (OS) was evident between the sTRA and non-sTRA groups of patients, but among the patients assigned to the non-sTRA group, those receiving adjuvant therapy had significantly longer overall survival (OS) than those who did not (p < 0.001, Log-rank test) (Figure 4B). Among the patients assigned to the sTRA group, no difference was observed. Both the TCGA and ICGC data sets showed this relationship. Other classifiers for the classical subtype gave similar results (Figure 4C), but not as consistently between data sets as the sTRA classifier.

In a parallel approach, we asked whether the directly measured amount of sTRA in primary tumors associated with a lack of response to adjuvant therapy. We used multimarker immunofluorescence^16, 25^ to quantify sTRA and CA19-9 in tissue microarrays (TMAs) that included tumors from patients who had long (> 3 years) or short (< 1 year) OS following surgery, and who either did or did not receive adjuvant therapy (Supplemental Table 2 and ref. ^26^). We quantified the markers using previously developed software that enables unbiased, automated quantification of multimarker IF data^16, 27^ (Figure 5A and Supplemental Table 2). The sTRA and CA19-9 levels showed little correlation with each other (Figure 5B), consistent with the two markers indicating different groups of tumors. The sTRA levels, but not the CA19-9 levels, were higher in the short-OS group (p = 0.0077, Wilcoxon Rank-Sum test) (Figure 5C). Likewise, in receiver-operator characteristic analysis to distinguish long from short OS, a test for area-under-the-curve (AUC) unequal to 0.5 was statistically significant using sTRA (p = 0.008, Wilcoxon test), but not using CA19-9 (Figure 5D).

**Figure 5.**
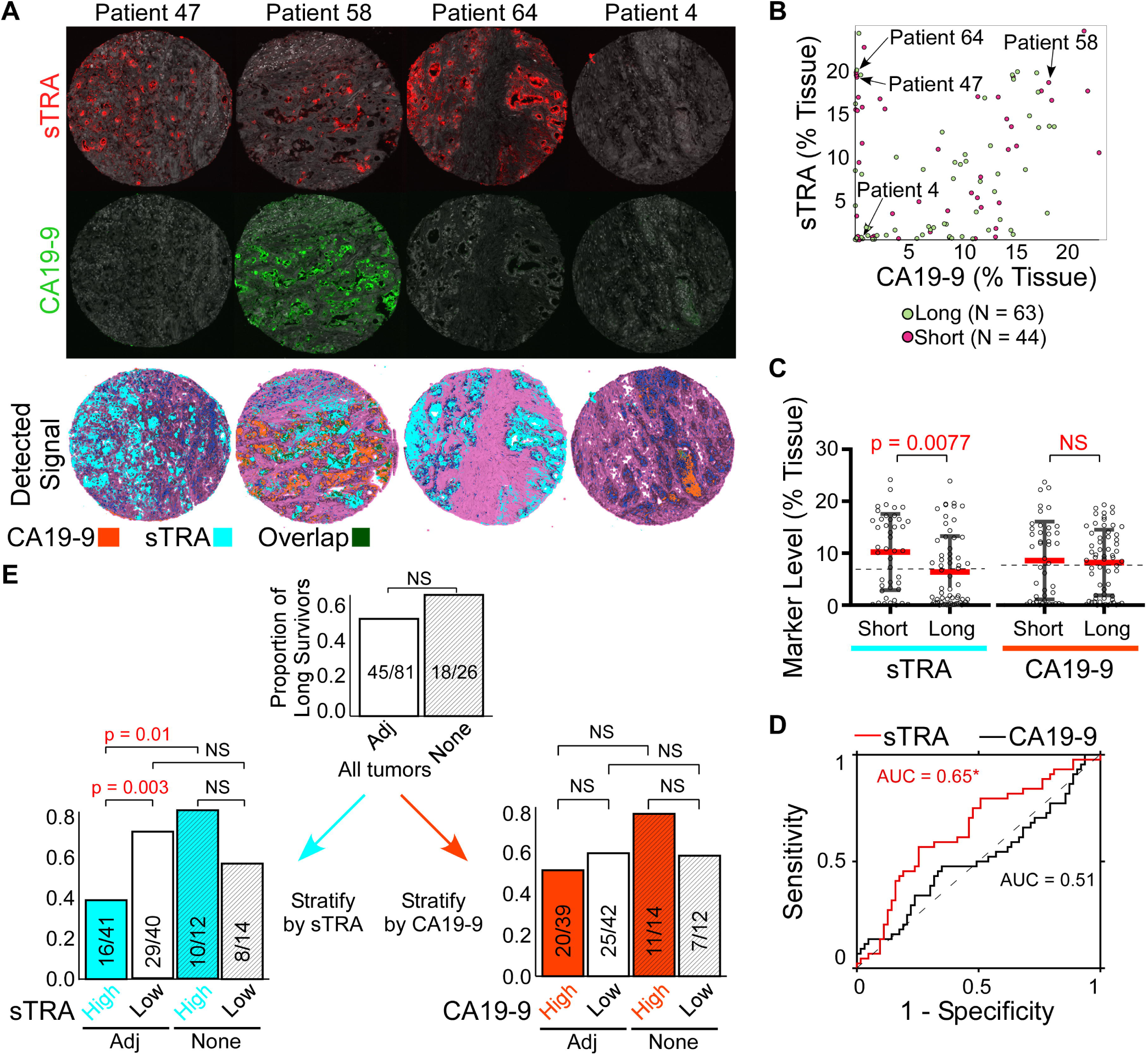
Tissue expression and outcomes. A) Raw immunofluorescence and detected signal of sTRA and CA19-9 in representative cores from the TMAs. Magnification is 4X. B) Comparison between the CA19-9 and sTRA levels in the patient tumors. Each point is the average of the three cores for a patient. C) Comparisons of sTRA and CA19-9 immunofluorescence data between the patient groups stratified by survival. The dashed lines indicate the median values that were used as cutoffs in panel E. D) Receiver-operator characteristic analysis for each marker to distinguish short survival (the cases) from long survival (the controls). AUC, area under the curve. E) Response to adjuvant therapy in subgroups. Adj. indicates the patients who received adjuvant chemotherapy following surgery. Using the median values of either marker given in panel C, the patients were further stratified either by sTRA (left) or CA19-9 (right).

The TMAs included similar proportions of long and short OS in the adjuvant and no-adjuvant groups (Figure 5E). Among the patients receiving adjuvant chemotherapy, those with high sTRA showed a significantly lower proportion of long OS than those with low sTRA (p = 0.003, Fisher’s Exact test). Among the patients with high sTRA, those receiving neoadjuvant chemotherapy had a significantly lower proportion of long OS than those not receiving neoadjuvant chemotherapy (p = 0.01, Fisher’s Exact test). No other comparison showed a significant difference. In addition, the Breslow-Day test for homogeneity of odds ratios (for association between survival and therapy) across biomarker-defined subgroups was highly significant for sTRA (p = 0.006) but was not significant for CA19-9 (p = 0.18). These results indicate a differential effect of adjuvant therapy on patient survival between the sTRA-high and sTRA-low groups.

Another set of TMAs contained samples from tumors that had been exposed to neoadjuvant therapy^16^. Although these TMAs did not reveal significant associations between survival and the individual marker amounts, possibly owing to changes in total glycan levels induced by the chemotherapy, they did show relationships between survival and relative biomarker abundance. The tumors that were dominant in cells producing only sTRA or only CA19-9, or that contained both types of single-labeled cells, were in the short survival group, with few exceptions (Supplemental Figure 4 and Supplemental Table 2). This relationship showed a connection between poor outcome and the outgrowth following neoadjuvant therapy of sTRA-dominant or CA19-9-dominant clones.

### Predicting rapid relapse using a blood test

A biomarker that is measurable in blood samples would have higher clinical utility than one accessible only from tissue, given the difficulties in sampling tissue from the pancreas. We therefore examined whether secreted glycans could be used as indicators of cellular expression. We measured the soluble levels of sTRA glycan using sandwich immunoassays, in which we detect sTRA on the proteins captured by one of three different capture antibodies (Figure 6A). The agreement between the cell-surface expression and the amount of glycan in the conditioned medium was high, with 25 of the 27 cell lines matching between cell-surface glycan expression and detection in the conditioned medium (Supplemental Figure 5). The agreement was also good for the organoids: 23 of 27 matched between cell-surface and conditioned-medium amounts for both sTRA and CA19-9. Furthermore, in a previous study we found that the peripheral blood glycans correlated with tumor glycans for cell-line xenograft mouse models, patient-derived xenograft mouse models, and human PDAC patients^16^. Thus, the secreted levels and blood levels are good indicators of the tumor expression of the glycans.

**Figure 6.**
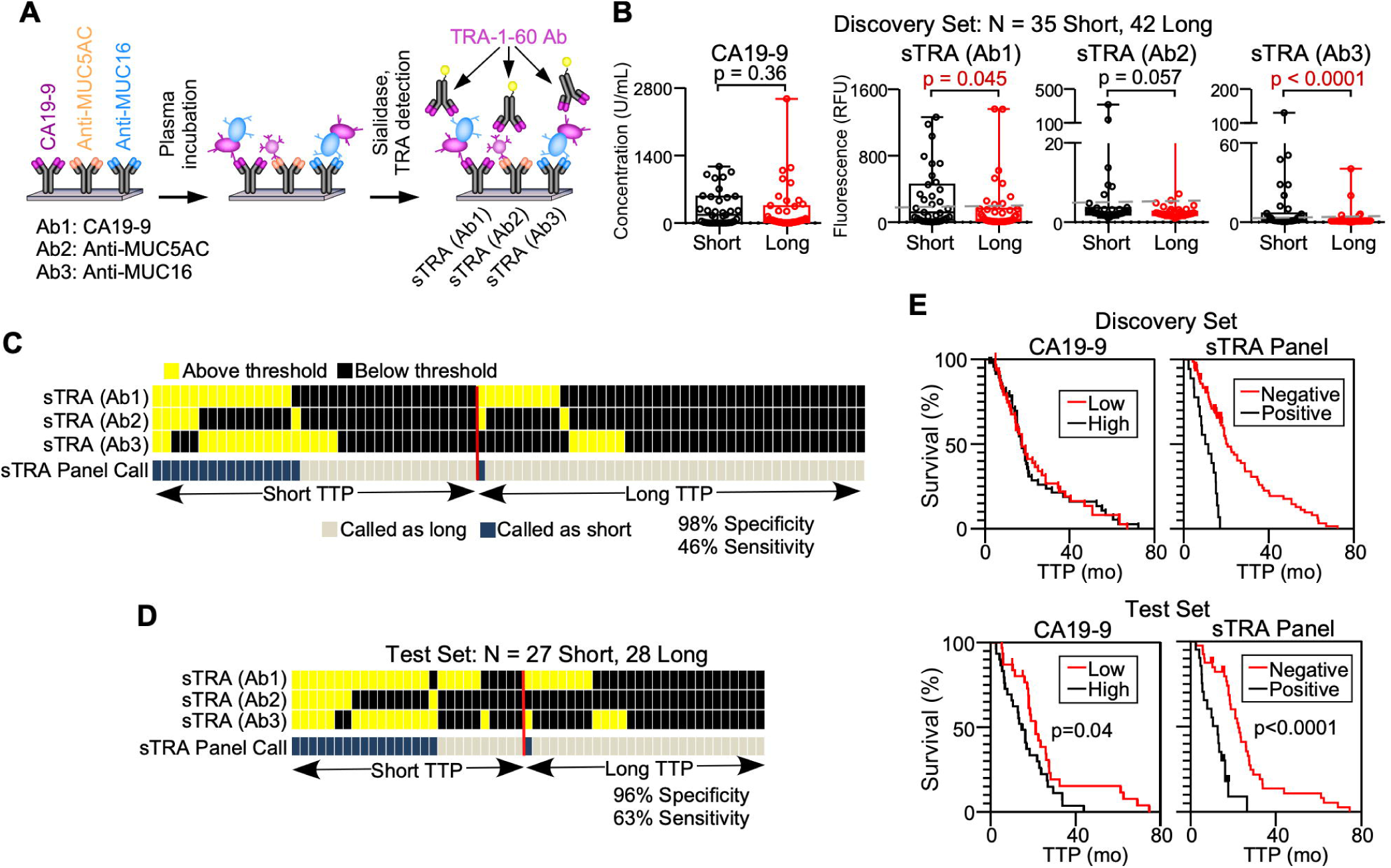
A blood test to predict rapid relapse following chemotherapy. A) Immunoassays used for measuring sTRA in patient plasma. B) Comparisons between short and long TTP, based on 18 months, of the indicated immunoassays. The values are the averages of three independent experiments. C) Patterns of high and low values across the three sTRA immunoassays in the discovery set. Each column represents a patient sample. Samples with elevation in two or three markers were classified as cases. D) Patterns of high and low values across the three sTRA immunoassays in the test set, using optimized thresholds. E) Survival curves in the discovery and test sets. For CA19-9, patients above the median were classified as high. For the panel, patients with elevations in two or three markers were classified as positive.

This relationship, combined with the above findings from cell culture and primary tumors, presented the possibility that high plasma sTRA identifies PDACs that do not benefit from systemic chemotherapy. We investigated this possibility among subjects who were beginning neoadjuvant therapy. Such a cohort would best reveal differences in resistance to chemotherapy, while avoiding potentially confounding major variations in the extent of disease or treatment history.

We measured sTRA and CA19-9 in plasma samples that had been acquired prior to the start of neoadjuvant therapy, first in a discovery set and then in a test set (data in Supplemental Table 3). We asked whether any of the biomarkers could serve as an indicator of short time-to-progression (TTP), as defined by radiographic evidence of progression. We dichotomized the patients using a cutoff of 18 months from the time of diagnosis, based on the approximate rate of 50% recurrence within one year after the completion of treatments.

In the discovery set, two of the sTRA immunoassays were significantly higher in subjects with short TTP than in subjects with long TTP (Figure 6B). CA19-9 did not show an association with TTP, nor did age or sex of the patients (Supplemental Table 3). We used a multi-marker classifier system called MSS^28^ to develop a three-marker panel. Patients elevated in two or more of the sTRA assays (using thresholds optimized for each marker, Figure 6B) were especially likely to have short TTP, as 16 of 17 had short TTP (94% PPV) (Figure 6C and Table 1). This result translated to 98% specificity (41/42 with long TTP were high in 1 or less) and 46% sensitivity (16/35 with short TTP were high in 2 or more). For CA19-9, a threshold that gave 95% specificity gave only about 3% sensitivity. To minimize the effect of overfitting on the estimate of panel performance, we further assessed the panel performance using five-fold cross-validation with 200-fold bootstrapping (re-samplings of the cohort). The improvements in cross-validated sensitivity and the average of sensitivity and specificity were statistically significant (p < 0.0001, Wald test based on bootstrap SE estimate) (Table 1).

**Table 1.** Performance of the sTRA Panel and CA19-9 in the discovery and test sets. CI, confidence interval. Bolded text of numbers indicates statistical significance. **p < 0.0001, *p < 0.01 (Wald test with bootstrap standard error estimate).

| Sample set/performance | Marker | Sensitivity (95% CI) | Specificity (95% CI) | (Sens + Spec)/2 (95% CI) |
| --- | --- | --- | --- | --- |
| Discovery Set |  |  |  |  |
| Naïve Performance | sTRA Panel | 45.7 | 97.6 | 71.7 |
|  | CA19-9 | 2.9 | 95.2 | 49 |
| Cross-Validated Performance | sTRA Panel | 45.4 (28.2, 63.7) | 92.5 (80.7, 97.3) | 68.9 (57.7, 78.3) |
|  | CA19-9 | 3.0 (0.1, 48.9) | 96.2 (85.9, 99.1) | 49.6 (44.8, 54.4) |
|  | Difference | **\*\*42.3 (21.7, 62.9)** | -3.7 (-11.8, 4.4) | **\*\*19.3 (8.1, 30.6)** |
| Test Set |  |  |  |  |
| Blinded Performance | sTRA Panel | 55.6 (37.2, 72.5) | 96.4 (78.1, 99.5) | 76.0 (64.8, 84.5) |
|  | CA19-9 | 11.1 (3.6, 29.2) | 96.4 (78.8, 99.5) | 53.8 (46.9, 84.5) |
|  | Difference | **\*\*44.4 (25.8, 63.1)** | 0 (-10.1, 10.1) | **\*\*22.2 (11.6, 32.9)** |
| Naïve Performance | sTRA Panel | 63 | 96.4 | 79.7 |
|  | CA19-9 | 11.1 | 96.4 | 53.8 |
| Cross-Validated Performance | sTRA Panel | 66.2 (44.8, 82.5) | 90.6 (75.4, 96.8) | 78.4 (64.6, 87.8) |
|  | CA19-9 | 13.3 (1.4, 61.6) | 96.0 (82.4, 99.2) | 54.6 (41.7, 67.0) |
|  | Difference | **\*\*52.9 (24.4, 81.4)** | -5.4 (-16.1, 5.3) | **\*23.8 (8.5, 39.0)** |

This is a provisional result, because the panel was optimized based on the data from the discovery set. We then applied the panel to a blinded test set. We applied the thresholds that were derived from the discovery set to the test set and made case/control calls on the blinded samples. The calls were sent to the collaborators who collected the samples, and upon comparison with the true outcomes data, the result was 96% specificity (16/35 with short TTP were high in 2 or more) and 56% sensitivity (15/27 with long TTP were high in 1 or less) (Table 1). The improvements in sensitivity and the average of sensitivity and specificity were statistically significant (p < 0.0001, Wald test based on 1000-fold bootstrap SE estimate, Table 1). We tested adjustments to the individual marker thresholds, considering that the thresholds were trained on a relatively small cohort, and we found that the optimized thresholds (naïve performance) gave 94% PPV (17/18 with two or more marker elevations had short TTP), 96% specificity (27/28 with long TTP), and 63% sensitivity (17/27 with short TTP) (Figure 6D and Table 1). Bootstrap analysis with cross validation confirmed the statistical significance of the improvement (Table 1). The three individual sTRA assays showed strong associations with short TTP (p = 0.008 to 0.00008, Mann-Whitney U test), as did CA19-9 (p = 0.007) (Supplemental Figure 6).

The sTRA panel also identified differences in TTP in Kaplan-Meier analysis (Figure 6E). Kaplan-Meier analysis is appropriate here because the cohort is a random selection of the patients seen in the clinic. In both sets, patients positive in the panel (elevated in 2 or more of the sTRA assays) had shorter TTP than the rest of the patients. In the test set, the difference was highly significant (p < 0.0001, log-rank test) for sTRA and moderately associated (p = 0.04) for CA19-9 (significance not evaluated in the discovery set).

To further test these associations, we obtained outcomes for a subset of patients in a previous study of these markers^17^ who had received neoadjuvant therapy (data in Supplemental Table 3). The study was not designed for this question but nevertheless could provide insights. In two separate cohorts, two of the sTRA assays trended with short overall survival (p = 0.05 and 0.06, Supplemental Figure 6). CA19-9 showed no such trend in either cohort. These findings substantiate the use of a blood test for sTRA to identify a subtype of pancreatic cancers that is resistant to chemotherapy.

## Discussion

This research demonstrates the use of the sTRA glycan to identify the PDAC cases that are highly resistant to chemotherapy. We used both cellular and blood-based immunofluorescence assays, as well as a surrogate gene-expression signature, to identify resistant PDAC. The identification was consistent across sample types—cell-culture models, isogenic cell lines, primary tumors, and blood plasma samples—and it was consistent in both the adjuvant and neoadjuvant settings. The immediate implication of this result relates to the development of treatment plans for patients with resistant PDAC. For patients with resectable PDAC, potentially morbid operations could be avoided if rapid relapse following surgery could be predicted *a priori*. For patients with metastatic disease and patients undergoing neoadjuvant therapy, a practical biomarker could guide the choices and comparisons of the treatment options. For example, FOLFIRINOX is suggested to be slightly better than gemcitabine for the classical subtype^10, 29^, possibly indicating a difference between the sTRA-positive and sTRA-negative types. Patient stratification also could improve the selection of patients that receive nab-paclitaxel^5^. Furthermore, the sTRA assay could be used in subgroup analyses of the many human trials currently underway, given that many trials do not meet primary objectives but show evidence of efficacy in subgroups. Trials could involve targeted therapies suggested from studies on the cell-culture and organoid models that are available for sTRA-positive and sTRA-negative PDACs. Thus, our biomarker has value for patient stratification using current options, as well as for research using model systems and in biomarker-guided drug trials.

A blood test has particular value in the clinical setting because physicians could stratify patients prior to any treatments, without a biopsy. A blood test also could extend the analyses of biomarker-based stratification beyond only the patients who have resected tumors or biopsy samples available. Furthermore, a blood test would capture secretions from the whole tumor rather than just the cells that are sampled by a biopsy, which may not reflect the heterogeneity of the tumor. Various blood tests have potential value for detecting or diagnosing PDAC, including mutated DNA in the circulation^30, 31^, tumor exosomes, and metabolites^32-34^, but they do not predict therapeutic responses. Highly elevated CA19-9 in the blood and the failure to drop to normal levels following neoadjuvant therapy or surgery^35, 36^ are unfavorable prognostic factors^37^, but CA19-9 does not indicate a distinct subtype or predict resistance to chemotherapy^35^. The sTRA blood test, in these initial studies, performed at a level that warrants further investigation as a biomarker to select for neoadjuvant or adjuvant chemotherapy. Blinded, prospective studies using a clinical assay will be required to fully assess the value of this test to patients and physicians.

If tissue is available from biopsy or resection, several cellular markers in addition to sTRA have the potential to be predictive. For example, high levels of the human equilibrative nucleoside transporter 1 (hENT1) showed a relationship with longer survival among patients receiving adjuvant gemcitabine^38^, presumably based on its role in the transport of gemcitabine and other drugs into cells. Immunohistochemical detection of KRT81 and HNF1A correlated with the basal and exocrine subtypes, respectively, and CYP3A5 indicated the exocrine subtype^21^. The analysis of immunohistochemistry data from TMAs indicated that high KRT81 corresponds to worse prognosis than high HNF81^29^, in line with the associations made by gene-expression profiling. GATA6 is an indicator of the classical subtype^7, 8^ and has been detected by in-situ hybridization in biopsy specimens^10^. Using IHC detection of GATA6 in TMAs, researchers found that tumors with high GATA6 had better prognoses than those with low GATA6, and that GATA6 had a weak association with resistance to 5FU^39^. These markers show promise, but their value for patient stratification will need to be validated in prospective studies.

This work extends previous findings relating to the prognostic and predictive value of the molecular subtypes. The classical subtype in previous research indicated a better prognosis than the basal subtype, but, on the other hand, it tended to benefit less from chemotherapy^9, 12^. This result is also consistent with an *in vitro* study in cultured cells, which indicated that classical-subtype cells were more resistant to gemcitabine^8^. But the relative value for prognosis versus treatment prediction was not clear, and a recent study suggested that the classical subtype is more sensitive to chemotherapy than the basal subtype^11^. The differences between the studies may result from several sources. The latter study included non-resectable, advanced PDAC (the COMPASS Trial), while the former studies involved resectable PDAC. Advanced cancer could be less responsive in general than localized disease. The COMPASS study also used a different classifier that was more stringent than the original, and the study was not designed to distinguish prognostic from predictive value, since it did not include a non-treatment control group. Overall, the results indicate that native prognosis and sensitivity to chemotherapy are not necessarily linked, and that the classical subtype is suboptimal for decoupling these traits.

The sTRA subtype seems to distinguish the traits better: it was indicative of chemotherapy resistance, but not of a poor prognosis. It encompassed resistant cancers of both subtypes and was more consistent than the classical/basal system in identifying resistance in the primary tumors. A valid model is that the sTRA subtype more precisely identifies resistant tumors, but the classical subtype could be more effective for stratifying by native prognosis. Ultimately, the typing of cancers and prediction of drug responses could involve both glycans and other types of markers. Further studies should focus on clarifying additional markers that are suggestive of other subtypes. The genes HNF1A, CDH17, LGALS4, and CYP3A5 have been variously assigned as markers of non-basal subtypes including exocrine and classical but without good agreement between studies^40^. The elevation of these genes in most sTRA cancers could indicate that a third subtype is at least partially encompassed by sTRA.

The sources of treatment resistance in the sTRA-positive cancers potentially could be discerned from the model systems and gene-expression studies, along with directions to test for new options. The sTRA cells could derive from a stem-cell population, given the structural similarity between the sTRA glycan and the glycan marker of iPSCs, as well as the stem-like gene expression. The development of resistant clones following therapy also is thought to arise from clonal diversity within tumors^41^, and stem-like cancer cells could be a minority subpopulation in early stages but become dominant in resistant cancers. Such a trajectory could underlie the development of tumors that are dominant in sTRA (Figure 5). Future studies could track changes in glycan markers in relation to clonal diversity.

Building on these findings, our next steps will involve the validation of clinical assays for prospective studies and the analyses of model systems in order to understand the susceptibilities of sTRA-positive cancers. Based on the gene-expression results, a successful path may involve metabolic approaches^42^. Alternatively, probing the sTRA-positive subtype for dependencies on particular nutrient sources may be feasible. These directions in research are made possible because we now have a practical assay to detect chemoresistant PDAC using either tissue or plasma. The use of such an assay in model systems, and then in clinical specimens to detect and follow the resistant subtype, should help both the development and the application of new treatments against PDAC.

## Acknowledgements

We thank the core services at the Van Andel Institute for expert support of this research, particularly the Optical Imaging, Bioinformatics and Biostatistics, Genomics, and Pathology and Biorepository cores. We thank Zachary Klamer, MS, at the Van Andel Institute for support in the processing and analysis of the plasma biomarker data; Toshinori Hinoue, PhD, at the Van Andel Institute for advising on the analysis of TCGA and ICGC datasets, and Christine Decapite at the University of Pittsburgh Medical Center for assistance with data coordination.

