## Supplemental data for "Detection of Chemotherapy-Resistant Pancreatic Cancer Using a Glycan Biomarker"

**Supplemental Tables**

- Supplemental Table 1. Gene expression analyses (Excel document)
- Supplemental Table 2. TMA data and analyses (Excel document)
- Supplemental Table 3. Plasma biomarker data (Excel document)
- Supplemental Table 4. Antibody information

**Supplemental Figures**

- Supplemental Figure 1. Cell line morphologies
- Supplemental Figure 2. Dose-response curves and chemosensitivity in CA19-9-expressing cell lines
- Supplemental Figure 3. Drug resistant isogenic cell lines are sTRA positive.
- Supplemental Figure 4. Outcomes associated with tissue staining
- Supplemental Figure 5. Immunoassay correspondence between surface expression and secretions
- Supplemental Figure 6. Additional plasma biomarker analyses in neoadjuvant therapy

**Supplemental Methods**

- Immunofluorescence
- Immunoassays
- Construction and sequencing of directional total rna-seq libraries
- RNAseq data processing
- Targeted genome sequencing
- Infinium qc array
- Gene-expression classification and survival analysis

The RNAseq data for 27 samples and the expression counts matrices have been submitted to NCBI GEO, **Series GSE146722**.

The release date is April 2021, or upon publication.

### Supplemental Tables

#### Supplemental Table 4. Antibody information.

| Antibody | Host | Clone | Target | Source | Cat. no. | Class |
|---|---|---|---|---|---|---|
| goat anti-mouse IgG-HRP | Goat | Polyclonal | Mouse IgG | Santa Cruz Biotechnology | SC2005 | IgG |
| goat anti-Rabbit IgG-HRP | Goat | Polyclonal | Rabbit IgG | Santa Cruz Biotechnology | SC2004 | IgG |
| CA19-9 | Mouse | 9L426 | Sialyl Lewis A | US Biologicals | C0075-03A | IgG |
| TRA-1-60 | Mouse | TRA-1-60 | Terminal *N*-acetyl-lactosamine, type 1 | Novus Biologicals | NB100-730 | IgM |
| MUC5AC | Mouse | 45M1 | Lewis B blood group antigen | Abcam | ab212636 | IgG |
| MUC16 | Mouse | X325 | Lewis blood group antigen | Abcam | ab10033 | IgG |

### Supplemental Figures

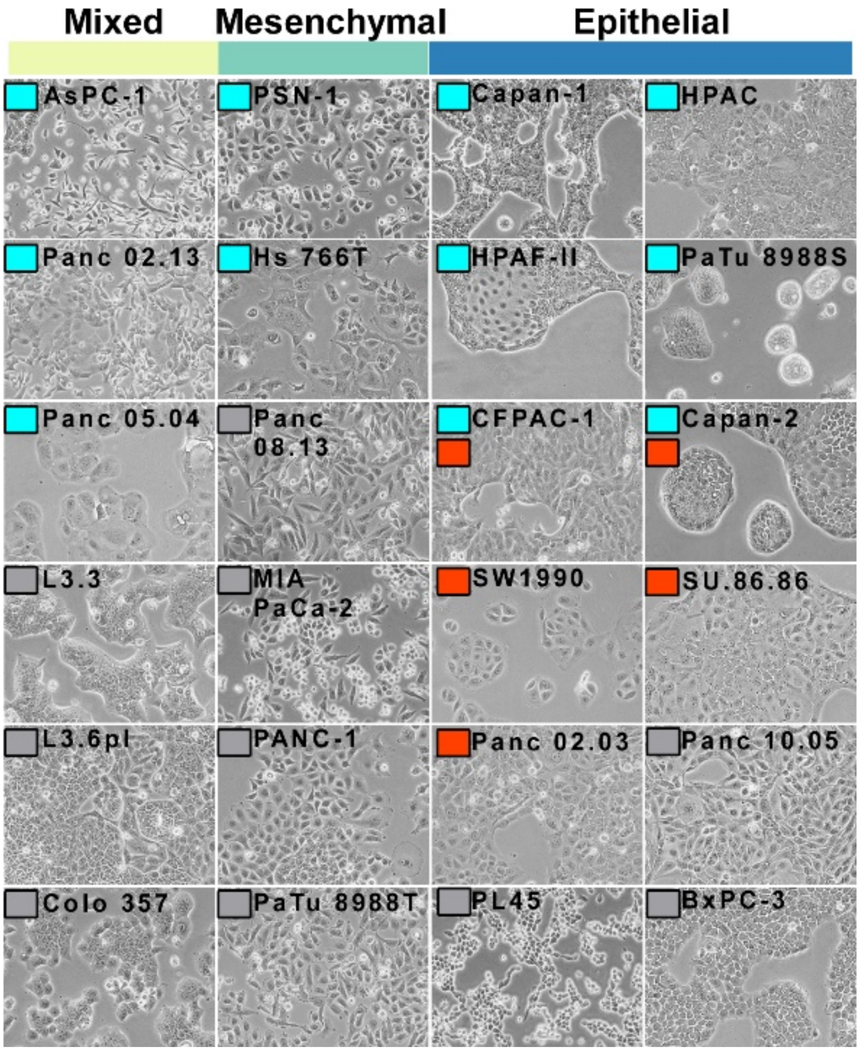


**Supplemental Figure 1. Cell line morphologies.** Brightfield images showing the morphologies of the cell lines. Magnification is 10X.

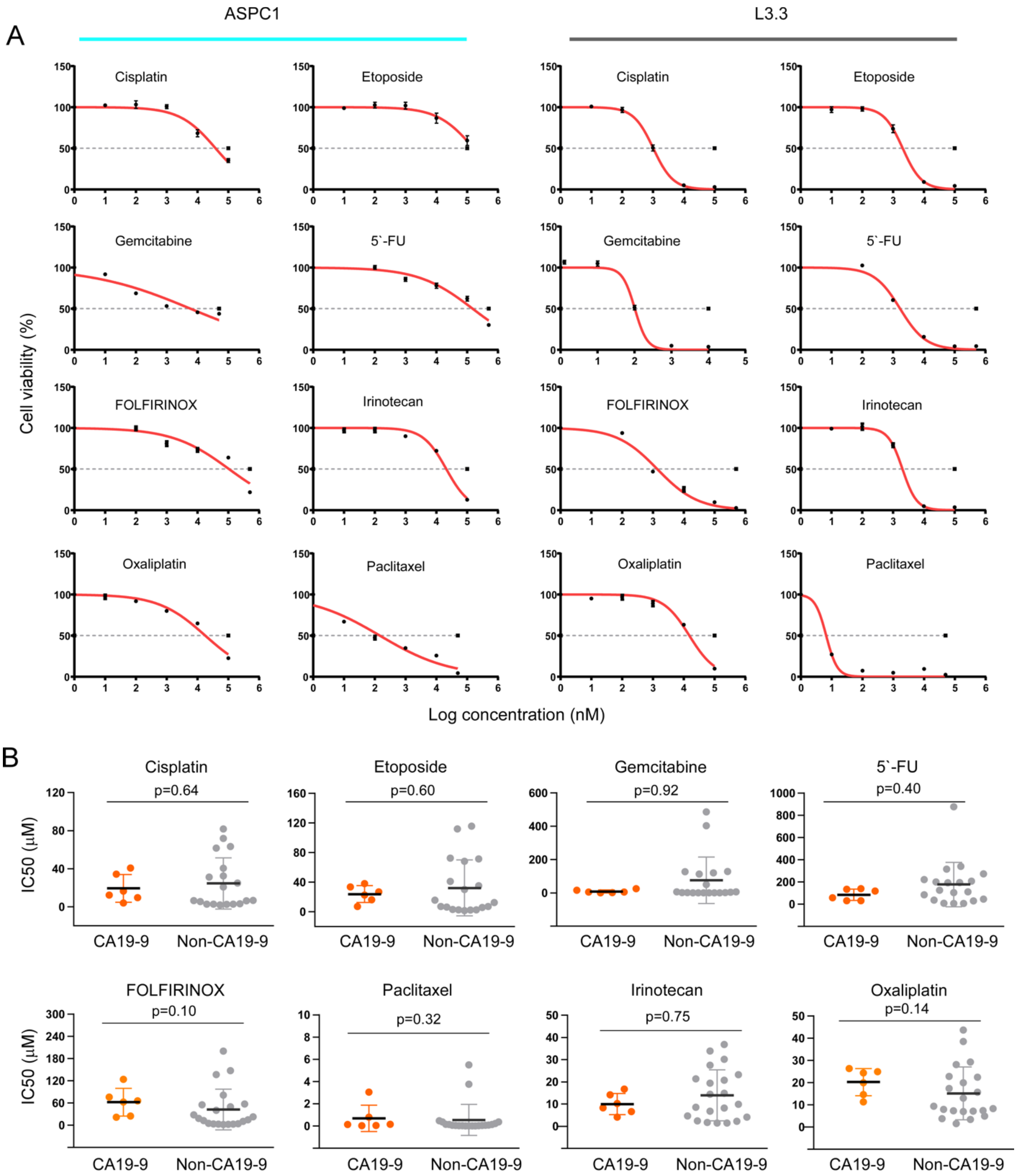


**Supplemental Figure 2. Dose-response curves and chemosensitivity in CA19-9-expressing cell lines.** A) Representative dose-response curves for cell lines. Each cell line was incubated with the indicated drug, titrated over a range of concentrations. Cell viability was measured by Cell Titer-Glo at 72 h, and dose-response curves were plotted to determine the $IC_{50}$ value for each combination. B). The $IC_{50}$ values calculated from dose-response curves, grouped by marker group. The p values are based on the Mann–Whitney test.

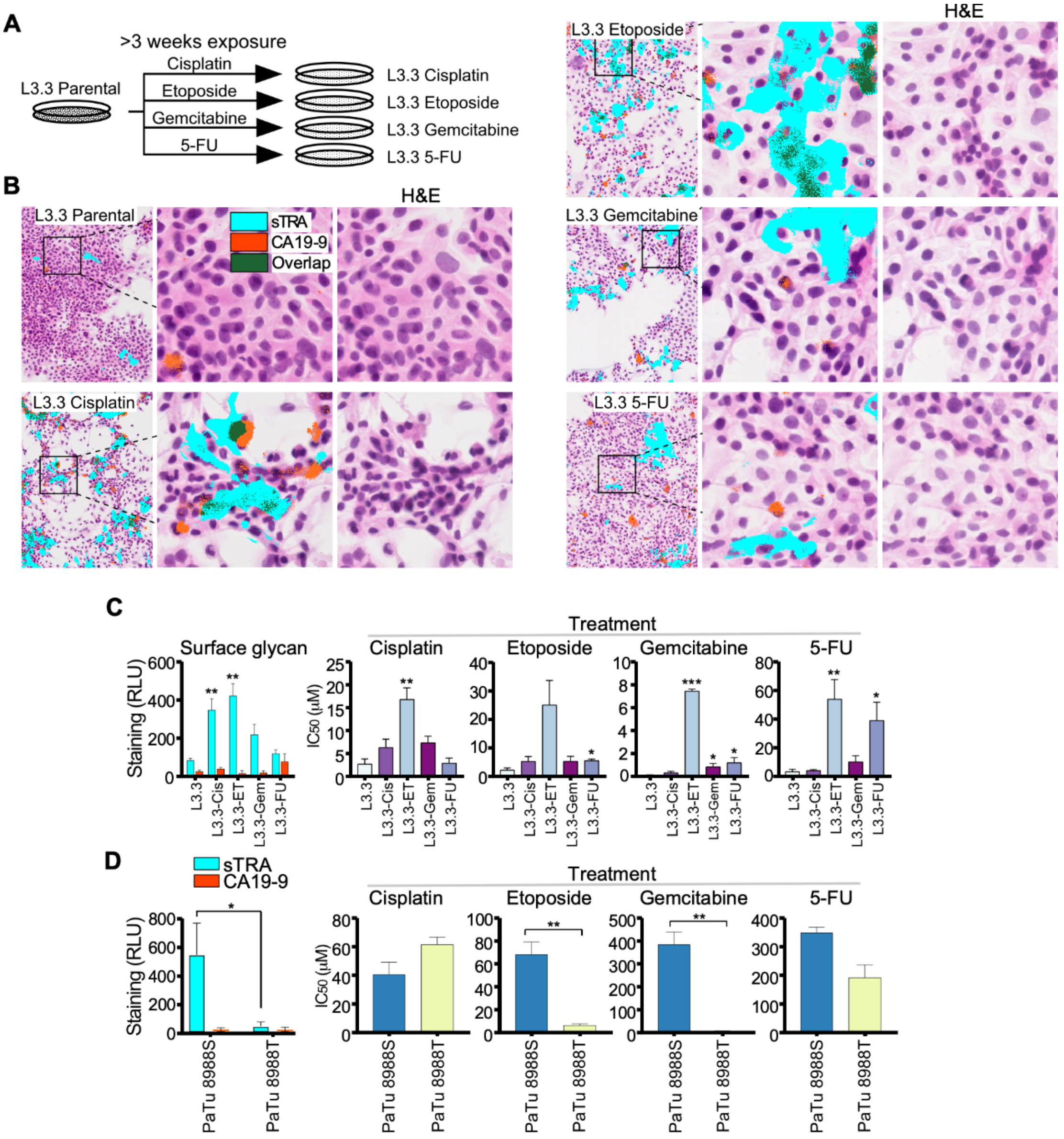


**Supplemental Figure 3. Isogenic cell lines.** A) Development of drug-resistant sublines of L3.3. B) Immunofluorescence of sTRA and CA19-9 in the parental and sublines. Magnification is 20X. C) Comparisons of the indicated features in the parental L3.3 cell line and the sublines. D) Comparisons between the PaTu8988S and PaTu8988T cell lines. The error bars are the standard error over 3 independent replicate experiments. H&E, hematoxylin and eosin. *indicates $p < 0.05$, **indicates $p < 0.01$. RLU, relative light units.

**A**

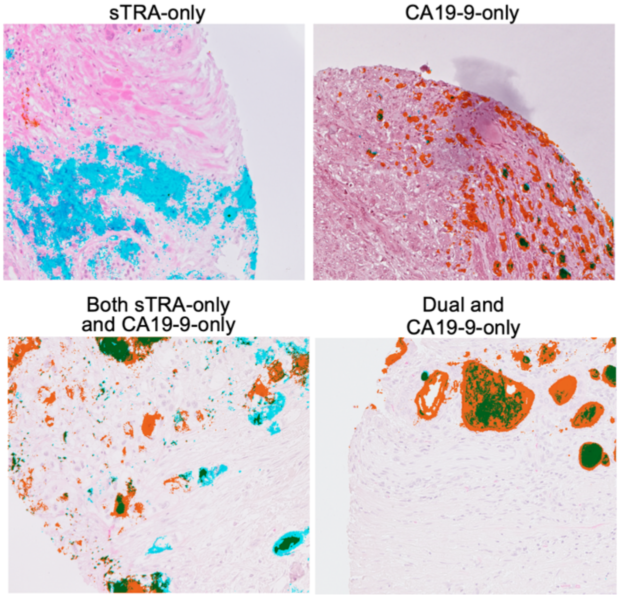


**B**

| UPMC | OS-Short | OS-Long |
|---|---|---|
| sTRA-only | 6 | 1 |
| CA19-9-only | 10 | 0 |
| Both | 4 | 0 |
| Dual-only | 6 | 18 |
| Low All | 19 | 17 |

**Supplemental Figure 4. Outcomes associated with tissue staining.** For each core on the TMAs, the SignalFinder program determined the percentage of the tissue that was positive for either sTRA or CA19-9. A) Representative immunofluorescence data from the UPMC TMAs. The detected signal from SignalFinder is overlaid on the H&E images. B) SignalFinder quantified the amount of staining in each core that consisted of only sTRA, only CA19-9, or overlapping signal (referred to as dual). These values were averaged over the three cores per tumor. Each tumor was classified into one of the five indicated categories based on the relative amounts of the types of staining. The full data and quantifications are provided in Table S2.

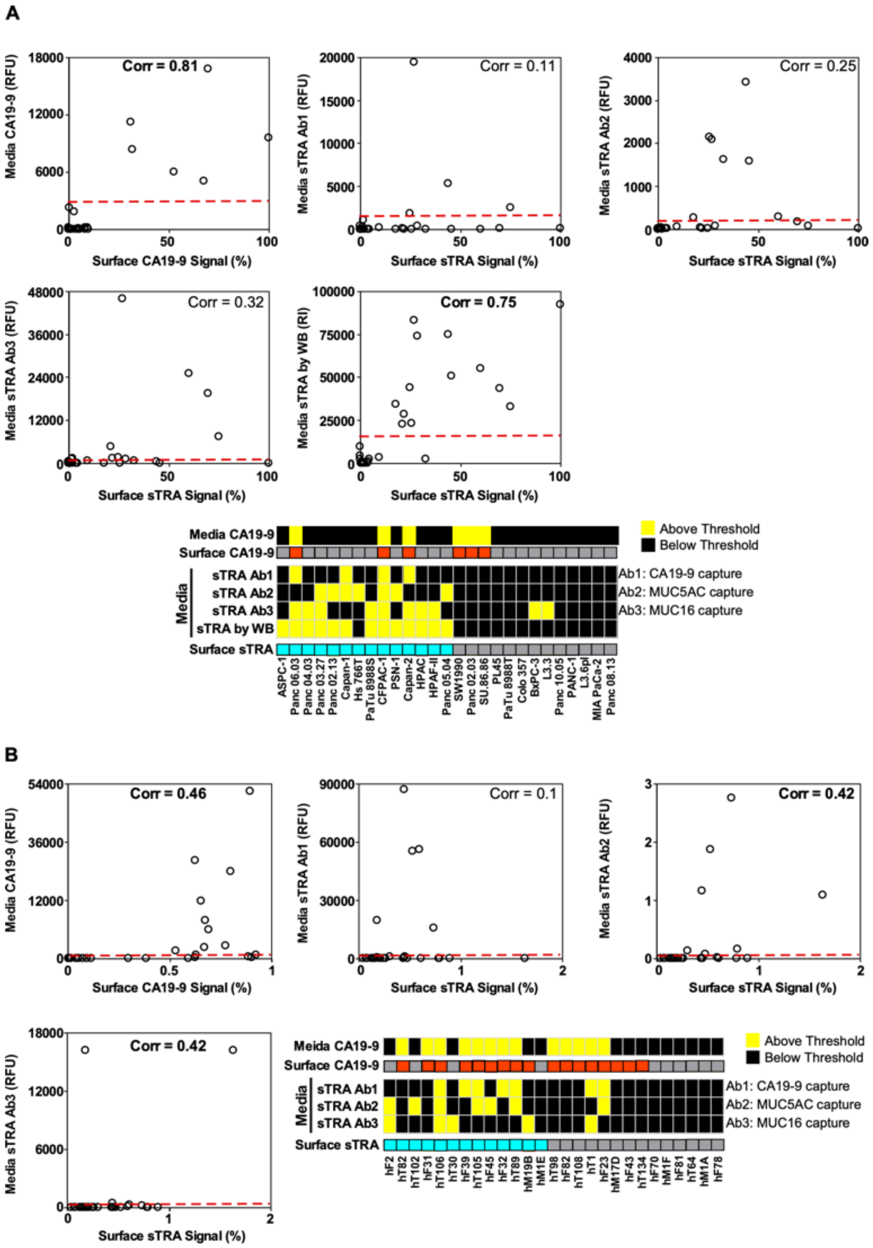


**Supplemental Figure 5. Immunoassay correspondence between surface expression and secretions.** A) Scatter plots of CA19-9 and sTRA between cell surface and secretions over 27 2D cell lines. The Pearson correlation coefficient was used to calculate the extent of correlation with significant correlation numbers in red. The matrix shows the patterns of high and low values across all immunoassays using secreted media. B) Scatter plots of CA19-9 and sTRA between cell surface and secretions over 27 organoids.

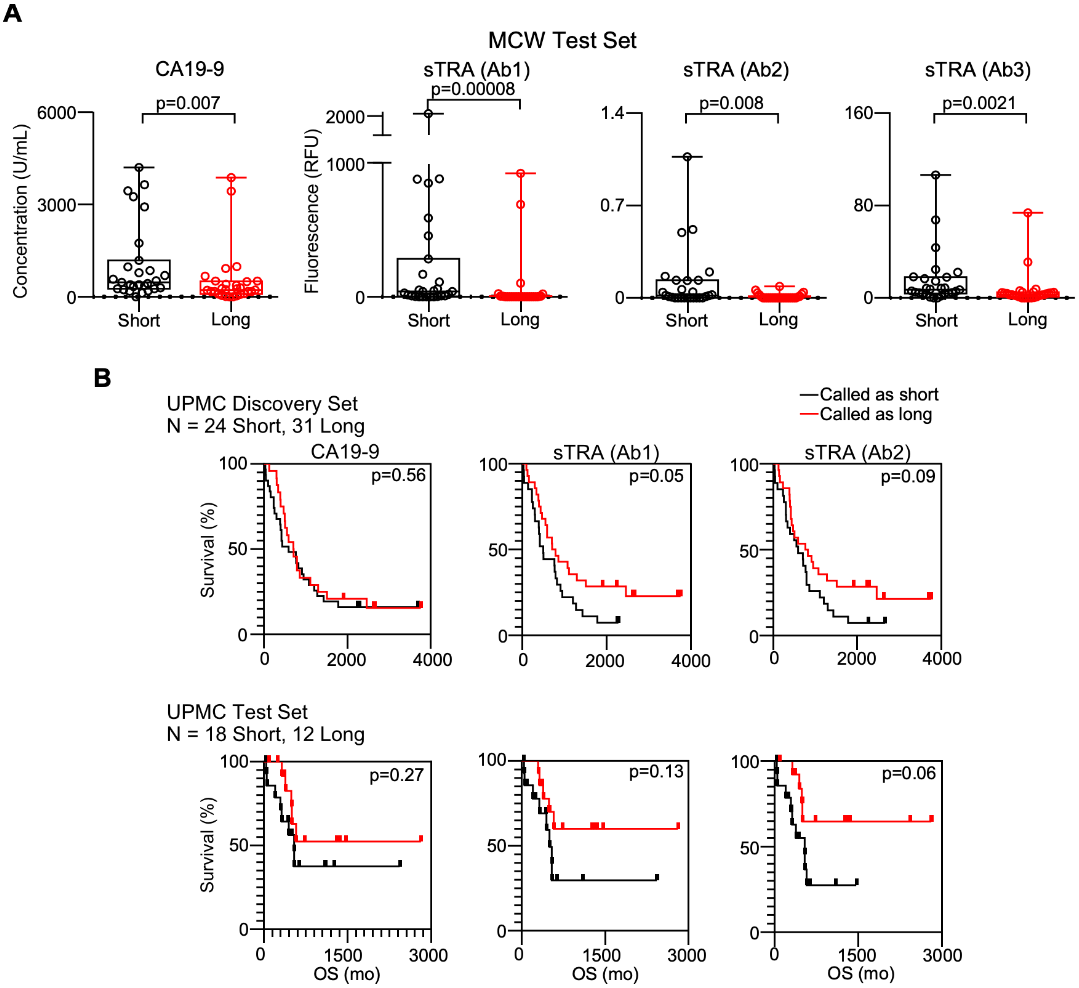


**Supplemental Figure 6. Additional plasma biomarker analyses in neoadjuvant therapy.** A) Individual markers in the MCW Test Set. B) Survival curves in retrospective analyses. The data are from previously published cohorts[1] from UPMC for three immunoassays, CA19-9, sTRA (Ab1), and sTRA (Ab2). Ab1 and Ab2 refer to the capture antibodies used in the sandwich assay in combination with sTRA detection (see Fig. 6, main text). Ab1 is the CA19-9 antibody, and Ab2 is anti-MUC5AC.

### Supplemental Methods

#### Immunofluorescence

All immunofluorescent and chemical stains were performed using 5-μm-thick sections cut from formalin-fixed, paraffin-embedded blocks. Paraffin was removed from the sections using CitriSolv Hybrid (04355121, Fisher Scientific), and tissue was rehydrated through an ethanol gradient. Following rehydration, antigen retrieval was achieved through incubating slides in citrate buffer at 100 °C for 20 min. Slides were blocked in phosphate-buffered saline with 0.05% Tween-20 (PBST0.05) and 3% bovine serum albumin (BSA) for 1 h at RT. Primary antibodies (Supplemental Table 4) were labeled for immunofluorescent staining with either Sulfo-Cyanine5 NHS ester (13320, Lumiprobe) or Sulfo-Cyanine3 NHS ester (11320, Lumiprobe) so that two primary antibodies could be used simultaneously in each round of staining. Dialysis was performed following labeling to remove unreacted conjugate and the primary antibodies were then diluted into the same solution of PBST0.05 with 3% BSA to a final concentration of 10 µg/mL. Slides were incubated overnight with this solution at 4 °C in a humidified chamber.

The following day, the antibody solution was decanted, and the slides were washed twice in PBST0.05 and once in 1X PBS, each time for 3 min. The slides were dried via blotting and then incubated with DAPI at 10 µg/mL in 1X PBS for 15 min at room temperature (RT). Two 5-min washes were performed in 1X PBS, and then slides were cover-slipped and scanned using a fluorescent microscope. All slides were scanned for fluorescence using either Vectra (PerkinElmer) for the TMAs or the Axio Scan.Z1 (Zeiss) for the organoid sections. Each system collected data for each field of view at three different emission spectra. All image data were quantified using the SignalFinder-IF software[2].

Following scanning, slides were stored in a humidified chamber. Coverslips were removed for the subsequent rounds of staining by submerging the slide in deionized water at 37 °C until the coverslip floated free (between 30 and 60 min). Fluorescence was quenched between rounds by incubating the slides with 6% $H_2O_2$ in 250 mM sodium bicarbonate (pH 9.5-10) twice for 20 min at RT. Subsequent incubations and scanning steps were repeated as described above with different primary antibodies.

To perform sialidase treatment, slides were incubated with a 1:200 dilution (from a 50,000 U/mL stock) of α2-3,6,8 neuraminidase (P0720L, New England Biolabs) in 1X reaction buffer (5 mM $CaCl_2$, 50 mM sodium acetate, pH 5.5) overnight at 37 °C. Slides were washed as described above, followed by subsequent antibody detections. Hematoxylin and eosin staining was performed following a standard protocol.

#### Immunoassays

The immunoassays were based on the method presented earlier[1]. The capture antibodies were CA19-9, anti-MUC5AC, and anti-MUC16, and the biotinylated primary antibodies were CA19-9 or TRA-1-60 (details in Supplemental Table 4). The secondary detection agent was Cy5-conjugated streptavidin (Roche Applied Science). We diluted the samples of human plasma (8-fold and 32-fold) and cell line or organoid media (2-fold and 8-fold), into a buffer (1X PBS with 0.1% Tween-20, 0.1% Brij-35, species-specific blocking antibodies, and protease inhibitor) and incubated each sample on an antibody array overnight at 4 °C.

For sTRA detection, an extra step of enzyme treatment before antibody detection was needed. After sample incubation we prepared α2-3 neuraminidase (P0728L, New England Biolabs, Ipswich, MA) at a concentration of 250 U/mL in the supplied reaction buffer and incubated on arrays overnight at 37 °C. We incubated each array with a biotinylated antibody (3 µg/mL in 1X

PBS with 0.1% Tween-20 and 0.1% BSA) and subsequently with Cy5-conjugated streptavidin (43-4316, Invitrogen, Carlsbad, CA) (2 µg/mL in the same buffer as the primary antibody). The slides were scanned for fluorescence at 635 nm using a microarray scanner (Innopsys InnoScan 1100 AL). The quantification of fluorescence was performed using SignalFinder-MA [2]. All plasma and media samples were repeated in at least three independent experiments. The CA19-9 values were obtained through the clinical laboratory services at the Medical College of Wisconsin for the discovery set, and they were determined using a custom immunoassay previously validated in the Haab laboratory[1] for the test set.

#### Construction and sequencing of directional total rna-seq libraries

Libraries were prepared by the Van Andel Genomics Core from 500 ng of total RNA using the KAPA RNA HyperPrep Kit with RiboseErase (v1.16) (Kapa Biosystems, Wilmington, MA USA). RNA was sheared to 300-400 bp. Prior to PCR amplification, cDNA fragments were ligated to Bioo Scientific NEXTflex Adapters (Bioo Scientific, Austin, TX, USA). The quality and quantity of the finished libraries were assessed using a combination of Agilent DNA High Sensitivity chip (Agilent Technologies, Inc.), QuantiFluor dsDNA System (Promega Corp., Madison, WI, USA), and Kapa Illumina Library Quantification qPCR assays (Kapa Biosystems). Individually indexed libraries were pooled and 75-bp, paired-end sequencing was performed on an Illumina NextSeq 500 sequencer using a 150-bp HO sequencing kit (v2) (Illumina Inc., San Diego, CA, USA), with all libraries sequenced to a minimum of 40 million reads. Base calling was done by Illumina NextSeq Control Software (NCS) v2.0 and output of NCS was demultiplexed and converted to FastQ format with Illumina Bcl2fastq v1.9.0.

#### RNAseq data processing

RNAseq libraries were generated using the KAPA RNA HyperPrep kit with RiboErase. Libraries were sequenced paired-end for 75 cycles on two Illumina NextSeq flowcells. Following demultiplexing, adapters and low-quality bases were trimmed using Trimgalore v0.4.2 (http://www.bioinformatics.babraham.ac.uk/projects/trim_galore/). Trimmed data was quality controlled with FastQC v0.11.7 and then mapped with STAR v2.5.2b to the hg38 genome using the default settings. Raw gene counts (mean of 27M/sample) generated by STAR were imported into R v3.6.0. Genes with greater than 10 counts in 2 or more samples were retained and the rest removed in order to minimize multiple testing adjustments and remove genes that are unlikely to give meaningful information. A quasi-likelihood negative binomial generalized log-linear model was then fit to the filtered count data using the weighted trimmed mean of M-values to normalize for library size and composition biases.

For all differential expression contrasts, the gene set was further filtered so that, for the subset of samples being used in the comparison, a minimum of two samples have more than zero counts. A quasi-likelihood, negative binomial, generalized log-linear model was then fit to the filtered count data using the weighted trimmed mean of M-values to normalize for library size and composition biases. GSEA v3.0 was used to test for enrichment of various gene sets in glycan phenotype groups. Significance of enrichment was tested using 1000 phenotype permutations. All gene sets were run simultaneously to adjust for multiple testing. Heatmaps were made using the R package pheatmap v1.0.12. Samples and genes were clustered using the default method of the pheatmap function. Normalized counts were used, median-centered across genes.

#### Targeted genome sequencing

Libraries were prepared by the Van Andel Genomics Core from 50 ng of high-molecular-weight genomic DNA using the KAPA Hyper Prep Kit (v5.16) (Kapa Biosystems, Wilmington, MA USA). In brief, DNA is sheared to an average size of 200 bp, then end-repaired and A-tailed DNA fragments were ligated to to Bioo Scientific NEXTflex Adapters (Bioo Scientific, Austin, TX, USA). Libraries were amplified and pooled for targeted capture in batches of 16 libraries, using 31 ng of each indexed library. Capture baits were designed using IDT xGen Lockdown Probes and Reagents (Integrated DNA Technologies, Coralville, IA, USA) with minimal modification. Quality and quantity of the captured library pools were assessed using a combination of Agilent DNA High Sensitivity chip (Agilent Technologies, Inc.), QuantiFluor dsDNA System (Promega Corp., Madison, WI, USA), and Kapa Illumina Library Quantification qPCR assays (Kapa Biosystems). A 150-bp, paired-end sequencing was performed on an Illumina MiSeq sequencer (Illumina Inc., San Diego, CA, USA) at the Michigan State University RTSF Genomics Core. Base calling was done by Illumina RTA3 and output of NCS was demultiplexed and converted to FastQ format with Illumina Bcl2fastq v1.9.0.

For all data, adapters and low-quality sequences (median $Q<20$) were removed with Trimgalore. Data was then mapped to hg38 using bwa mem. Following mapping, the median non-duplicate read depth for captured regions was 156x. From this high-depth data, variants were called using the best-practice workflow of the GATK4 tool Mutect2. Variants were then annotated using the command-line variant effect predictor tool from Ensembl using release 92. The annotated variant files were converted to mutation annotation format (MAF) using vcf2maf (https://github.com/mskcc/vcf2maf), and oncoplots and lollipop plots were generated using the R package maftools v1.6.15. To confirm specific mutations, eight samples were also re-sequenced paired-end for 300 cycles on an Illumina MiSeq.

#### Infinium QC array

DNA was quantified by Qubit fluorometry (Life Technologies) and 200 ng of DNA from each sample was processed by the VARI Genomics Core using the Illumina Infinium QC array v1.0 (Illumina), which contains some 16,000 SNPs focused on sex determination, ethnic ancestry, ADME, and genetic linkage markers. DNA was amplified, hybridized to the QC Array bead chip, and an extension reaction was performed using flurophore labeled nucleotides per the manufacturer’s protocol. Array beadchips were scanned on the Illumina iScan platform and probe specific calls were made using Illumina Genome Studio software.

#### Gene-expression classification and survival analysis

The gene classifier for sTRA included the top 14 up-regulated and top 14 down-regulated genes ($p < 0.02$ after multiple-testing correction) in sTRA-expressing cells versus sTRA-negative cells in the RNAseq analysis of 27 cell lines. We obtained the expression data (Z scores) for the gene list from The Cancer Genome Atlas (TCGA dataset provisions, https://www.cbioportal.org/)) and the International Cancer Genome Consortium (ICGC data portal https://dcc.icgc.org/projects/PACA-CA). For each case of PDAC, we calculated a score by subtracting the average of the down-regulated genes from the average of the up-regulated genes. The patients above the median score were classified as sTRA-signature positive. To classify patients as classical or basal in the ICGC data, we used the same algorithm but with a previously-identified gene list that included 23 classical-associated gene and 23 basal-associated genes[3]. The Kaplan-Meier survival curves were generated with GraphPad Prism 6. The subjects who were alive at the end of the study or who died of other causes were censored.
